# Population genomic structure of sorghum landraces across landscape, environment and culture

**DOI:** 10.1101/2025.11.02.686151

**Authors:** Eleanna E. Vasquez Cerda, Emily S. Bellis, Aayudh Das, Emma R. Slayton, Geoffrey P. Morris, Jesse R. Lasky

## Abstract

The spread of staple crops to diverse environments over time and their current genetic structure may reflect historical dispersal by humans, sustained human preference for particular traits, and adaptation to local environments. Sorghum is a drought-tolerant crop native to Africa cultivated by hundreds of millions of smallholders globally. Here we examined the ecological context of population-genomic structure of 1,806 sorghum landraces across Africa and Eurasia to infer the relative contribution of environmental and cultural factors to sorghum genetic diversity across different relative time periods. Sorghum landraces were spatially and linguistically structured at a large-scale and within subregions, following a pattern of isolation by distance. Within regions, much of genomic structure was best explained by a mechanistic model of human travel time. In our assessment of hierarchical linguistic structure, we found that language families explain 4% of genomic variation while individual languages explain 13% of genomic variation, suggesting the importance of human culture and relationships in gene flow and selection. Variance partitioning showed that travel time, language, and climate explain up to 27% of genomic variation among landraces. We also observed regional differences in the degree of genetic relatedness across space and time in our assessment of shared ancestry. East Africa showed particularly strong geographic turnover in genomic composition and haplotype sharing, while West Africa showed substantial haplotype sharing even over large distances, signifying some rapidly spreading lineages. Thus, space, travel time, and culture likely capture important forces controlling sorghum genomic variation, but these factors operate heterogeneously over space.

## Introduction

The rate of gene flow and geographic patterns of relatedness may be shaped by dispersal ability, geographic distance and population size (Bohonak, 1999). In natural plant populations, genetic material like seeds and pollen are dispersed through vectors like wind or animals (Ellstrand, 2014; Rousset, 2000) and the gene flow distribution has a higher density of events within short distances and a lower density at long distances (Ellstrand, 2014; Ibrahim et al., 1996; Rousset, 1998; Slatkin, 1993). With limited dispersal and no selection, a pattern of isolation by distance may be observed; more distant populations may have increased genetic differentiation due to reduced gene flow and drift (Kimura & Weiss, 1964; Sexton et al., 2014; Wright, 1943). Among other natural forms of dispersal, human-mediated dispersal can play a role in gene flow dynamics, particularly in long-distance dispersal events (>5 km) (Auffret et al., 2014; Bullock et al., 2018; Ellen & Platten, 2011; Wichmann et al., 2008). Human-mediated dispersal events may be accidental (Wichmann et al., 2008), or intentional through the exchange of seeds among communities (Menamo et al., 2021) and human migration (Boivin et al., 2017; Ellen & Platten, 2011).

By modeling human-mediated dispersal for crops, inferences can be made about plant gene flow dynamics and spatial connectivity (Gutaker et al. 2020). Human movement can be modeled by identifying least cost paths (LCP), which in this context assumes that humans traverse heterogeneous landscapes by considering routes that minimize energy expenditure (Adriaensen et al., 2003; Gowen & de Smet, 2020; Rees, 2004). Rates of human movement between points may be modeled using estimated travel time, where locations connected by shorter travel times are likely to have more frequent movement. If human movement is the primary vector of crop gene flow, then least cost travel times for humans may better predict the genetic differentiation between crop populations than geographic distance (Gutaker et al. 2020).

Humans also play an essential role in the distribution of crop genetic variation through intentional and unintentional selection. Breeders intentionally cross distinct genotypes to confer beneficial traits (Ohadi et al., 2017) like increased yield and stress tolerance (Swarup et al., 2021). For smallholders, selection may be driven by perceived habitat suitability (Atinkut & Mebrat, 2016; Seo & Mendelsohn, 2008). Similarly, culture may inform conscious or unconscious preference for varieties not explained by climate (Faye et al., 2019) such as non-adaptive traits like color (Lacy et al., 2006). Cultural factors like ethnicity, farmer social structure and language have been associated with population structure in crops like maize, barley and sorghum (Benz et al., 2007; Gilabert et al., 2025; Samberg et al., 2013; Westengen et al., 2014). This association may be explained by the ‘farming-language codispersal hypothesis,’ which holds that languages have historically dispersed with the diffusion of agriculture, aligning with migration and population expansion events (Diamond & Bellwood, 2003). Using language as a proxy for culture may help untangle the confounding relationships affecting genetic diversity such as spatial structuring and environment as well as environment and culture (Leclerc & d’Eeckenbrugge, 2012; Perales et al., 2005; Velásquez-Milla et al., 2011). By incorporating these factors into models of population structure, we may better understand the individual and conjoint effects of spatial connectivity, human movement, language and climate on sorghum genetic structure.

Patterns of genetic relatedness across space may be temporally heterogeneous, as demography and patterns of gene flow across landscapes can change over time. For example, changes over time in the slope of relatedness decay over distance have been observed in European humans (Al-Asadi et al., 2019; Ralph & Coop, 2013). While longitudinal genomic samples over time are often missing, identity by descent tracts of different lengths, which are shared portions of the genome passed down over generations from a common ancestor between two haplotypes, can reveal temporal dynamics in demographic history of populations (S. R. Browning & Browning, 2012). The length of identity by descent tracts (along the genetic map), can be utilized as a relative estimate of timescales where longer segments represent more recent genetic relatedness while shorter segments represent older relationships, allowing us to look back in time (B. L. Browning & Browning, 2013). Pairwise comparison of identity by descent tracts between samples may reveal temporal changes in the degree of shared ancestry and the relationship between genetic relatedness and geographic space (Al-Asadi et al., 2019; Ralph & Coop, 2013).

Sorghum is a self-pollinating drought-tolerant crop that is geographically widespread and adapted to a wide range of climatic conditions (Menamo et al., 2021; Morris et al., 2013; OECD, 2014; Olatoye et al., 2018). For millennia since its domestication in Africa, sorghum has served as a staple for people across Africa and Asia with versatile use as food, fiber and fuel (OECD, 2014; Venkateswaran et al., 2019). Researchers have postulated that sorghum originated in East Sudan and Western Ethiopia 4,600 to 10,000 years ago based on archaeological evidence (Kimber, 2000; O. Smith et al., 2019; Winchell et al., 2017) and more recently, based on spatially explicit genetic analyses (Gilabert et al., 2025). Cultivated sorghum likely diffused along ancient overland and sea trade routes within continental Africa and Asia (Kimber, 2000; Venkateswaran et al., 2019). For example, sorghum dispersal from Africa to the Sind-Punjab region and India may have occurred from ports in East Africa via the Indian ocean and along the Red Sea (Sabaean lane) around 3000 B.C and 2000 B.C respectively (Harlan & Stemler, 1976; Kimber, 2000). Within continental Africa, diffusion of varieties followed North-South and East-West axes around >3000 B.C and 2000 B.C respectively (Kimber, 2000). Putative dispersal routes of sorghum varieties (Kimber, 2000) may have been determined by landscape and environmental features favoring seed movement (i.e. the primary mechanism of gene flow in a self-pollinating species).

Culture is associated with sorghum genetic variation regionally (Faye et al., 2019) and across Africa (Gilabert et al., 2025). The effects of geography and climate on population structure have also been assessed at a large-scale (Lasky et al., 2015). However, the relative effects of language, human movement, geography, and environment have not been considered across global sorghum landraces. Gilabert et al. (2025) recently identified genetic clusters in sorghum that were associated with specific climates and language families, which when subdivided exhibited fine-scale population structure. For example, two genetic clusters in West Africa were associated with the Atlantic-Congo language family and Mande subclass. By conducting an effective migration surface analysis in Africa, both Gilabert et al. (2025) and Morris et al. (2025) found evidence of geographic structuring, specifically corridors across space with increased gene flow and areas with reduced connectivity such as the Ethiopian highlands.

However previous studies have not assessed the combined influence of language, geography, and climate over time across Africa and Eurasian sorghum.

We build upon findings in previous studies by estimating spatiotemporal and linguistic differences in genetic structure as well as the relative effect of spatial connectivity, human movement, language, and climate on sorghum genomic diversity. We estimate hierarchical genetic differentiation in 18 language families and 97 individual languages. By conducting a variance partitioning analysis, we disentangle the portion of genetic variation explained by language families relative to geographic distance, climate, and human movement. Here we highlight both geographic distance and human-induced dispersal patterns as distinct drivers of population structure using isolation by distance and travel time models across continents, regions and language families. To infer temporal differences in geographic structure we estimate the distribution of shared ancestry among paired landraces across space, which to date has not been implemented in sorghum landscape genetics. Here we ask:

1) Is sorghum genetic variation better explained by geographic distance or a human movement model? Dispersal limitation and genetic drift likely result in a positive linear relationship between genetic distance and geographic distance or travel time. However, travel time may be a better predictor of genetic diversity as geographic barriers that constrain human-mediated dispersal are considered.
2) How does the hierarchy of human language diversity explain sorghum genomic variation? Shared preferences for traits or similar agronomy and environment among individuals sharing the same language may act as selective forces that drive genetic differentiation between landraces associated with different language.
3) What is the relative contribution of geographic isolation, language and climate in explaining sorghum genomic variation? While climate may explain a portion of genomic diversity due to local adaptation to climate, we expect a large proportion of this relationship is collinear with geography due to spatial autocorrelation in climate gradients (Coelho et al., 2023; Ruiz Miñano et al., 2022; Sork et al., 2010). Collinearity between climate and language may reflect differential selection of locally suitable varieties and climatic barriers.
4) How do the above patterns change among regions and over time? Processes like spatiotemporal habitat heterogeneity may shift patterns of gene flow across space and over time. This may be reflected in differences in the spatial distribution of landraces across regions and time. If sorghum landraces follow a pattern of isolation by distance, we expect the number of shared identity by descent segments to decay with increasing geographic distance between paired landraces. Furthermore, if the processes we described above operate heterogeneously over time, we expect larger identity by descent tracts to display the strongest isolation by distance pattern as more recent relationships may be confined across geographic space (Ralph & Coop, 2013).

## Methods

### Genomic and Climatic Data

We obtained genotype by sequencing (GBS) data for 1,806 sorghum landraces collected across Africa and Eurasia (Figure 1) from the study by (Hu et al., 2019). For most of the analysis below, we excluded the relatively isolated northeast Chinese landraces, because their status as geographic outliers could overly influence some analysis. For wavelet analysis, which allows localized estimations of population structure, we included the Chinese landraces (see below).

**Figure 1.**
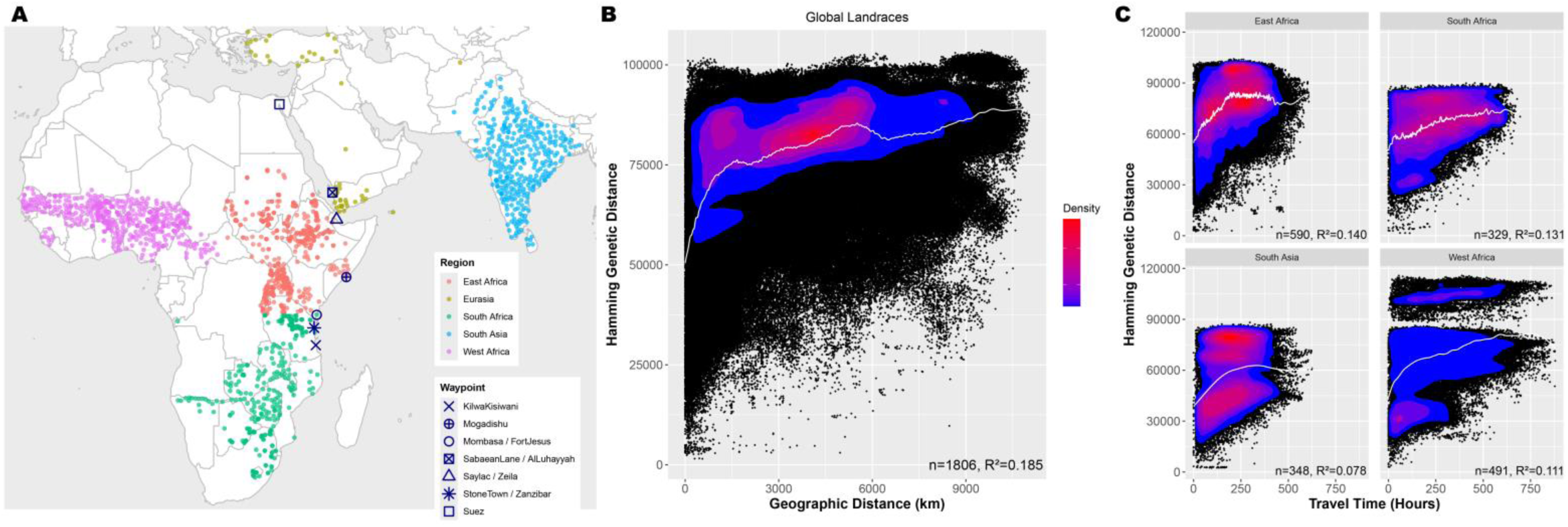
A) Map of 1,806 sorghum landraces used in regional analyses (East, West, and South Africa and South Asia) and Eurasia which was included in global analyses. The waypoints used to connect intercontinental landrace pairs are denoted by symbols. **B)** scatterplot of geographic and genetic distance globally, **C)** scatterplot of travel time in subregions East Africa, South Africa, South Asia, and West Africa. Spline (white line) added to show distribution pattern. Density bar indicates concentration of landraces (blue-low, red-high).

We considered temperature and moisture as variables due to their effects on crop performance and evidence that crops are adapted to these aspects of environment (Bannayan et al., 2011; Gutaker et al., 2020; Riha et al., 1996; Slingo et al., 2005; Sloat et al., 2020; Verón et al., 2015). We extracted climate variables for each georeferenced landrace using WorldClim v.2.1 data with 30 arcsecond resolution (Fick & Hijmans, 2017). For temperature we used 5 variables: mean temperature of wettest quarter, mean temperature driest quarter, temperature seasonality, mean temperature coldest quarter, max temperature of warmest month. For moisture we considered 6 variables: aridity index, precipitation of the driest quarter (log(x+1) transformed), precipitation of the driest month (log(x+1) transformed), precipitation seasonality, precipitation of wettest month and precipitation of the wettest quarter.

### SNP Filtering

To avoid likely read mapping errors, we removed SNPs with more than 5% heterozygotes in a custom script and converted to ped/map format using gdsfmt v.1.38.0 and SNPRelate v.1.36.0 in R v.4.3.3. We also removed rarer variants (MAF <= 0.05) using PLINK v.1.9.0-b.7.7 leaving a total of 135,085 SNPs.

### Genetic Distance

To consider the extent to which geographic distance and travel time influence genetic divergence between landraces, we estimated genetic distance for each pairwise landrace (Hamming distance, PLINK v.1.9.0-b.7.7). We found genetic distance was the best fit between genetic and geographic distance (R^2^ = 0.185) compared to Rousset’s (1/1-F_ST_) (R^2^=0.000), pairwise F_ST_ (R^2^=0.003), Nei’s (R^2^= 0.184), and Euclidean genetic distance (R^2^=0.1672) measures.

### Geographic Distance

To test for signatures of isolation by distance, we calculated geographic distance between pairs of landraces using the Vincenty ellipsoid, using the distm function from *geosphere* v.1.5-18 (Hijmans, 2024). This method follows an ‘as the crow flies’ route finding (i.e irrespective of barriers) for each pairwise landrace.

### Travel Time

As an attempt to more accurately estimate rates of sorghum gene flow between populations, we modeled human-mediated dispersal of sorghum landraces by calculating travel time of the least cost path (LCP) between landraces. To do so, we first modeled human movement across terrain of heterogenous slope, where steep slopes slow the speed of movement (Tobler, 1993). Speed as determined with Tobler’s hiking function assumes that the maximum walking speed of 6 km/hr is reached on a slight decline (-0.05° slope) while the speed of walking across flat land is 5 km/hr (Goodchild, 2020; Tobler, 1993). To obtain slopes we used elevation data from Worldclim v.1.4 with 30sec resolution. Over sea we simulated low-technology sailing (Gutaker et al., 2020). For travel over water, we estimated 3 nautical miles or 1.54 m/s) (Irwin et al., 1990; Slayton, 2018; Surface-Evans & White, 2012), causing travel across water to be 1.1x faster than across flat land (Gutaker et al., 2020).

For each pair of landraces, we estimated travel time by generating a conductance layer, which we use to estimate resistance across a given geographic range using gdistance v.1.2-2 (van Etten, 2017). The conductance layer contains values equal to the reciprocal of travel time between a focal cell and its neighboring 8 cells. Using the costDistance function from gdistance v.1.6.2 in R v.4.2.1, we calculated the number of hours for the LCP between each pair of landraces; this function computes the reciprocal of the conductance matrix values (1/conductance = distance/speed) between two sets of coordinates so that *travel time = speed/distance* for movement over land and sea (Gutaker et al., 2020; Slayton, 2018). Due to computational constraints to working with a single conductance layer across Africa and Eurasia at the desired resolution, we generated separate conductance layers for Africa and South Asia and then connected intercontinental landrace pairs via putative important waypoints. We included seven potential waypoints of ancient trade: Suez (Egypt), Mombasa/Fort Jesus (Kenya), Saylac/Zeila (Somalia), Kilwakisiwani (Tanzania), Sabaean Lane/Al Luḩayyah (Yemen), Stone Town/Zanzibar (Tanzania), and Mogadishu (Somalia) based on historical sorghum transportation or ancient trade routes (Harlan & Stemler, 1976; Murdock, 1959). For example, the Sabaean Lane in modern-day Yemen was an important overland and maritime connection between Africa and South Asia (Harlan & Stemler, 1976). To estimate travel times for pairs of African and Eurasian landraces, we first calculated the least cost path from all landraces within each continent to each waypoint. Then, for each unique landrace combination between different continents, we extracted the waypoint with the shortest combined travel time. To calculate travel time for distant landraces or computationally intensive regions within Africa, we generated additional conductance layers by constraining layers to the smallest size possible within 5° of the minimum and maximum longitude and latitude. We also calculated travel time for landraces with unique locations by rounding the coordinates to 0.25° and assigning the output to the remaining corresponding landraces.

### Spatial Model Selection

To assess which model (geographic distance or travel time) was a better predictor of genetic distance, we extracted AIC values from a linear mixed effects model where individual genotypes have random effects (Clarke et al., 2002) using the MLPE.lmm function from ResistanceGA v.4.2-10 (Peterman, 2018) in R v.4.2.1. We compared transformations of geographic distance, specifically the square root, log and untransformed version, and found that the untransformed version was the best fit to genetic distance based on AIC.

### Language Structure

To evaluate whether landraces genomic variation is linguistically structured, we first obtained language data from the glottolog database (Hammarström et al., 2022), which contains georeferenced language and language family data across the world. First, we assigned each landrace to a language and language family based on the geographically closest language, as geographic distance is correlated with linguistic typological distance (Auer et al., 2013). To estimate population structure among language families and languages for 135,085 SNPs, we first filtered for language families with more than five languages, leaving a total of 785 landraces. We then used the varcomp.glob function from hierfstat (Goudet, 2005) v. 0.5-11 to calculate hierarchical *F_ST_*. To estimate *F_ST_* among language families across all 1,806 landraces, we used Weir and Cockerham’s estimate from hierfstat.

### Regional Analyses

The processes that generate population genomic differences among landraces, such as human-mediated gene flow, may operate differently in different regions. Thus, we stratified our isolation by distance analyses by subregions based on putative dispersal patterns of sorghum varieties mentioned in (Kimber, 2000; OECD, 2014). We conducted SNP filtering and calculated genetic distance for each subregion in isolation. We then tested the relationship between geographic and genetic distance within each region to estimate geographic change in isolation by distance.

To further understand geographic patterns of population structure across spatial scales and locations, we implemented wavelet analysis of genome-wide landrace differences. Wavelet analysis allows estimation of spatial scale-specific genomic differences among genotypes and can be used to localize changes in these patterns across the landscape (Lasky et al., 2024). We calculated genome-wide wavelet dissimilarity at a range of spatial scales from ∼50 m to ∼8100 km for all landraces from Africa and Eurasia. We compared the observed genomic dissimilarity at each scale with a null distribution generated from permuting landrace locations (i.e. a null model of panmixia, used for comparison).

### Temporal changes in population genomic structure

While some models may present evidence of geographic structuring, it would be difficult to disentangle which genetic relationships are more recent across geographic space, particularly if the distribution of individuals or pattern of isolation by distance is expected to vary with time (Al-Asadi et al., 2019; Ralph & Coop, 2013). Temporal differences in geographic and genetic structure can be measured through comparison of identity by descent tracts, opening a potential window into past relatedness (Lawson et al., 2012). Genotype sharing of identity by descent segments across segments of different lengths can represent a range of timescales: smaller shared identity by descent segments representing older relationships, and larger shared segments arise from recent relationships.

To estimate identity by descent sharing, we first filtered SNPs and individuals in the VCF file to match other analyses using VCFtools (Danecek et al., 2011) v.0.1.16. The remaining SNPs we phased using BEAGLE v. 5.5 (27Feb25.75f) (B. L. Browning et al., 2021) and a sorghum genetic map from (Marla et al., 2025). To identify identity by descent segments across haplotypes between paired landraces, we used the software Refined IBD v. 17Jan20.102 (B. L. Browning & Browning, 2013) along with the phased data and genetic map. Following a script from (Al-Asadi et al., 2019), we created an identity by descent sharing matrix for different length bins (1-3 cM, 3-5 cM and >5 cM). We calculated the mean number of shared identity by descent segments across haplotype pairs for each paired landrace within each region. To assess geographic distance decay in the sharing of identity by descent segments, and regional differences, we tested a Poisson weighted regression model of geographic distance with region as an interaction versus pairwise identity by descent sharing, using regional sample sizes as weights (Ralph & Coop, 2013).

### Variance Partitioning

To assess the unique and redundant prediction of population genomic structure by language families, geographic distance, travel time, temperature and moisture, we conducted variance partitioning with redundancy analysis (RDA). RDA is an ordination of multivariate responses (SNPs) with multivariate predictors (e.g. language, climate) that allows estimation of the proportion of genetic variation explained by sets of factors. We reduced the dimensions for geographic distance and travel time matrices by filtering for unique coordinates and extracting a set of spatial eigenvectors with positive values, which we isolated with the pcnm function from vegan following (Bauman et al., 2018). This resulted in 158 and 421 eigenvectors for the geographic distance and travel time models respectively across 1,403 landraces. We used the varpart function from vegan to conduct variance partitioning (Oksanen et al., 2012) v.2.6-10 in R v.4.3.3.

Additionally, we estimated changes in the processes driving population structure over time by conducting distance-based variance partitioning for pairwise sharing of different identity by descent segment bins (1-3 cM, 3-5 cM and >5 cM). To reduce skew and convert the haplotype sharing matrix into a measure of genetic distance, we applied a sqrt((1/(1+x))^2) transformation, where x is the mean number of shared identity by descent segments for a pair of landraces. We then used this matrix in a distance-based RDA (Legendre & Anderson, 1999) and variance partitioning with the same covariates described above.

## Results

### Pattern of isolation by distance and travel time

In our analysis of pairwise geographic and genetic distance, we found strong evidence of isolation by distance in 1,806 sorghum landraces across Africa, Eurasia and South Asia (Figure 1A). We observed an increase in genetic distance as geographic distance increased (R^2^ = 0.185) (Figure 1B). This pattern indicates that landraces that are geographically proximate are more genetically similar than those that were further apart. Similarly, we observed a pattern of isolation by travel time, specifically an increase in genetic distance as travel time hours increased (R^2^ = 0.188, S1). Contrary to our expectations, AIC favored a geographic distance model (AIC = 2.1086e+07) over travel time (AIC = 2.1089e+07) globally, suggesting some limitations to our global travel time model.

### Regional differences in spatial structuring

In contrast to global models, we found that within regions, the travel time model was most favored. The travel time model was a better fit in East Africa (Geographic distance (G)=1447729, Travel time (T)=1447668, n=367), West Africa (G=1825684, T = 1825608, n=410) and South Asia (G=1209018, T=1208859, n=338) whereas in South Africa (G=650351, T=650371, n=329) the favored model was geographic distance. The difference in model selection results globally versus within regions may be due to differences in the magnitude of distances, and the domain over which our model worked best. Globally, 59% of pairwise distances were over 3000 km and traveled through our waypoints between continents, while regionally only 0.6% were over 3000 km. Additionally, our waypoints and intercontinental travel may have been erroneous.

We observed an approximately linear increase in genetic distance with increasing travel time hours and geographic distance within each region (Figure 1C & S2). However, while all regions follow a pattern of isolation by distance, the patterns of spatial structuring (i.e density and distribution of paired landraces across geographic space) appeared to be variable. Notably, landraces in West Africa contained the most genetically distinct landraces within short distances. The distinct division between two clusters and plateau in genetic distance in West Africa is indicative of highly diverged landraces and limited gene flow between the two groups. These genetic clusters correspond to the genetically divergent *guinea margaritiferum* variety (S5-7) and the more common, *guinea*, *durra* and *caudatum* type present in the region (Gilabert et al., 2025; Morris et al., 2013, 2025). We observed a similar pattern in South Asia, though to a lesser degree, indicating the divergence between the genetically distinct *guinea* and *durra* varieties that dominate opposite wet and dry sides of the Indian subcontinent. Regional analyses indicate that 11, 13 and 14% of genetic variation was explained by travel time constraints in West, South and East Africa respectively. On the other hand, only 7% was explained by the same model in South Asia, which suggests that the pattern of spatial structuring may be stronger within Africa (where sorghum is native) compared to South Asia.

There were also major differences revealed among regions in scale-specific wavelet analyses, that allowed us to identify particular locations of major genomic turnover (Figure 2). At the ∼387 km scale there were still a few regions with significantly low dissimilarity. In particular, Chinese landraces had significantly low dissimilarity at this scale, potentially reflecting their more recent colonization (<2 kya, versus e.g. ∼5 kya for colonization of Punjab) (Kimber, 2000; Qingshan & Dahlberg, 2001) and the rapid spread of closely related genotypes. By contrast, at the same 387 km scale there was particularly high dissimilarity along the Rift Valley in Ethiopia, a region of high sorghum diversity and great topographic and climate heterogeneity (Lasky et al., 2015), as well as in eastern India, a region where the very distinct *guinea* and *durra* landraces meet (Wang et al., 2020). At the scale of ∼1039 km, we found significantly high dissimilarity nearly everywhere, especially West Africa (Burkina Faso to Nigeria) and SE Africa (Zambia to Mozambique), which both correspond to the early axes of sorghum spread out of east Africa (Kimber, 2000) and are regions of turnover in major genetic clusters (Wang et al., 2020), as well as western India and Pakistan, which correspond to sharp rainfall gradients along which sorghum landraces may be locally adapted (Lasky et al. 2015).

**Figure 2.**
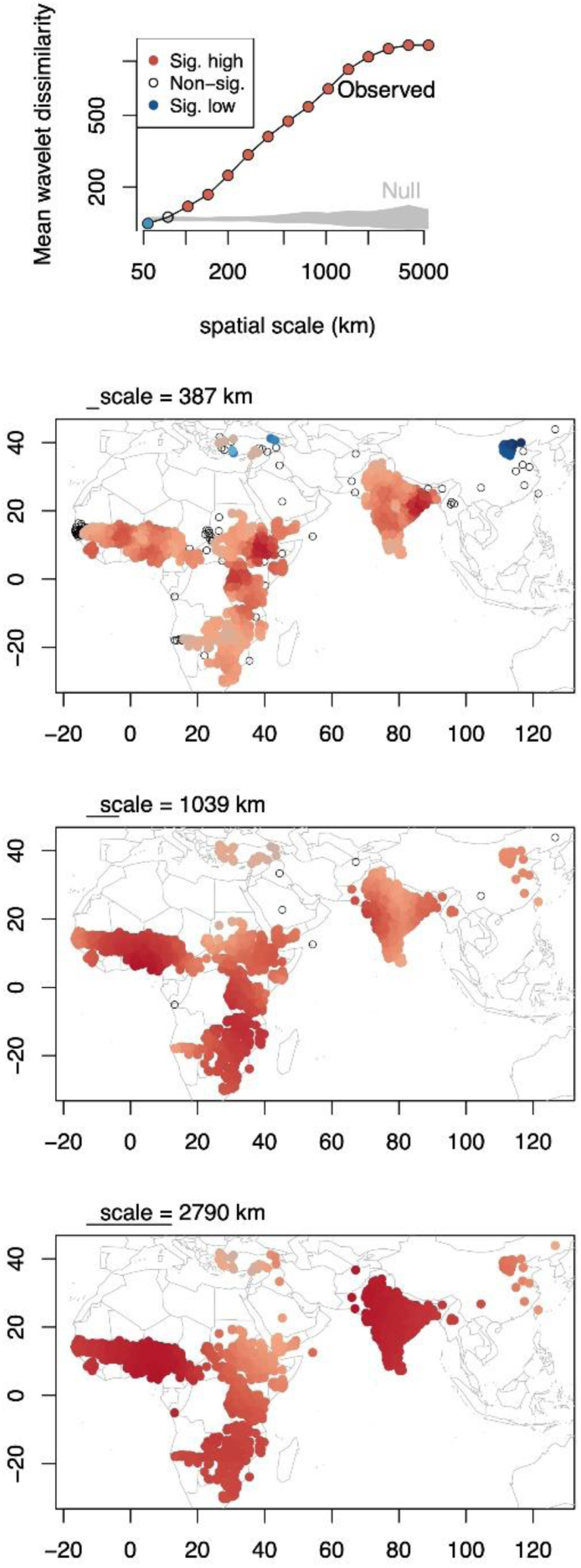
Scale-specific and localized population genomic structure in sorghum landraces identified using wavelet analyses, with blue indicating less and red more genomic turnover than panmixia, respectively. (top panel) The global mean wavelet dissimilarity increases with spatial scale, becoming greater than a null expectation from panmixia (grey) at ∼100 km. (bottom three panels). (bottom panels) Wavelet dissimilarity at a range of spatial scales, with significantly low dissimilarity (e.g. northeast Chinese landraces at 387 km scale) shown in blue and high dissimilarity shown in red, non-significant locations lack colors.

### Sorghum landraces are linguistically structured

In our analysis of language structure, we used hierarchical *F_ST_ (HF_ST_),* metrics to estimate the degree to which languages and language families explained population genetic structure in sorghum landraces in 785 sorghum landraces, (i.e. those landraces assigned to language families with at least 5 languages) and associated 18 language families. We found that 4% of genomic variation was among language families while 13% was among individual languages (Figure 3A). Additionally, 10% of genetic variation within language families was among languages. Our estimate of global *F_ST_* across 1,806 sorghum landraces and 42 associated language families similarly indicated that only 4% of genetic variation was explained by language families. When we stratified this analysis across continents, we found that 3% and 2% of genomic variation was explained by language families within Africa and South Asia respectively. In our estimate of *HF_ST_*, we observed weaker linguistic structuring in South Asia relative to Africa: in South Asia there was no genetic variation among language families while 8% of genetic variation was both within individual languages, and languages within language families. On the other hand, in Africa, 2% of genomic variation was among language families while 12% was within individual languages, and 10% of variation within language families was within languages. These findings indicate that sorghum landraces are linguistically structured, but the portion of genetic variation explained by individual languages is greater than that of language families at a large scale and across continents. Comparatively, the degree of linguistic structuring was stronger in Africa than in South Asia.

**Figure 3.**
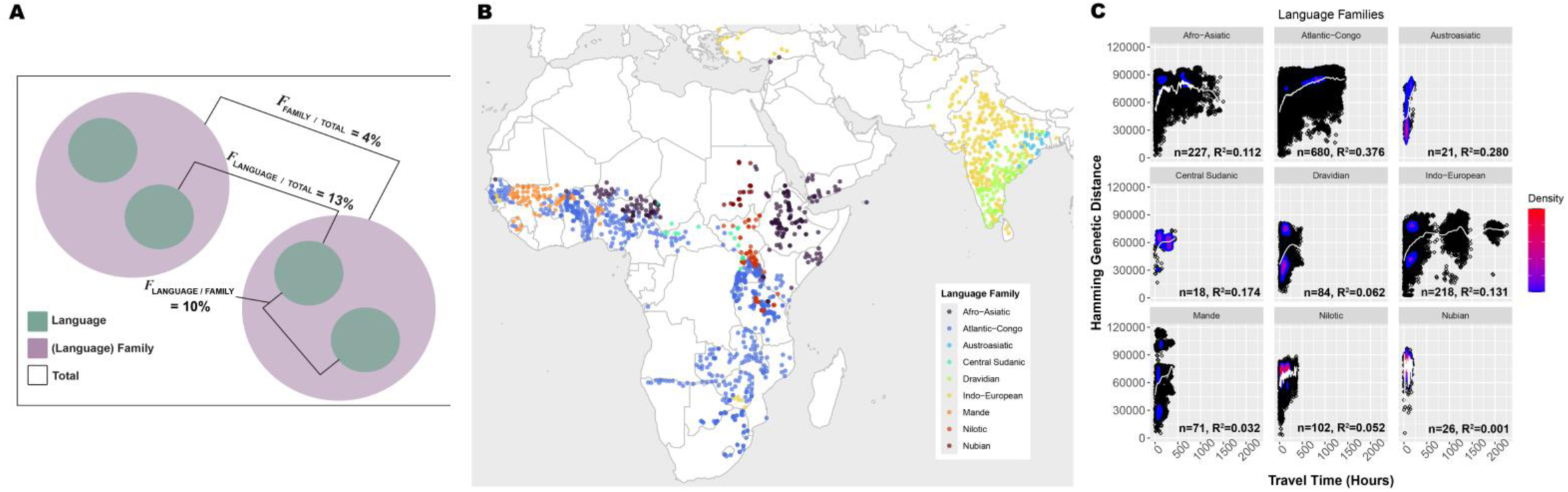
A) Diagram of population and subpopulation hierarchies and the associated HFst %. Out of 785 landraces, 13% of genetic variation is among language groups, 10% of variation within families is among individual languages and 4% is among language families. **B)** Map of language families observed in assessment of linguistic structuring. **C)** Scatterplot of geographic and genetic distance across individual language families. Spline (white line) added to show distribution pattern. Density bar indicates concentration of landraces (blue-low, red-high).

### Linguistic regions and isolation by distance

While the pattern of isolation by distance was present globally, this pattern varied across language families (Figure 3B & C). For example, although the Nubian and Austroasiatic language families had comparable sample sizes and geographic extents (Sudan and eastern India respectively), the linear models indicated that 0% and 28% of genetic distance was explained by geographic distance for each respective language, perhaps due to the role of large-scale rapid diffusion of sorghum along the Nile River in Sudan. Within the Atlantic-Congo language family, which had the largest sample size, 37% of genetic distance was explained by the travel time model while only 11% and 13% was explained by the model within the Afro-Asiatic and Indo-European language families respectively. Thus, among language families there was variation in patterns of isolation by distance. Although generally there was an increase in genetic distance as travel time increased across language families, the rate of increase in genetic distance was variable (slopes): in the Indo-European language family genetic distance steadily increased and began to plateau around 500 hours, while in the Afro-Asiatic language family the increase was nonmonotonic and steep with a decline around 500 hours. Conversely, within the Afro-Asiatic language family, which was comparatively geographically constrained, genetic dissimilarity increased rapidly as travel time increased and then decreased steadily after 500 hours. On the other hand, landraces within the Mande language family were predominantly constrained to West Africa yet exhibited evidence of genetic divergence at relatively short travel times. Just as in our regional analysis, this divergent genetic cluster may be associated with the genetically distinct *guinea margaritiferum* variety and the more common *durra* varieties. This finding follows the hypothesis of an independent domestication event within West Africa by individuals from the Mande linguistic group around 4500 B.C (Murdock, 1959). Linguistic differences in spatial structuring across the remaining language families may be attributable to variation in geographic extent within language families, regional differences in variety preference within language families, or variety-specific differences in adaptation to particular climates, which may be similar at extreme distances or highly variable within short distances/travel hours.

### Temporal differences in spatial population genomic structure

We used different identity by descent tract lengths to represent the degree of shared ancestry at different timescales where sharing of smaller (1-3 cM) and larger (>5 cM) represents older and more recent relationships, respectively. The mean tract length in our dataset was 3.9 cM, the median was 3 cM while the 1^st^ and 3^rd^ quantile were 2.1 cM and 5 cM respectively. We found the mean number of shared identity by descent segments across length bins decreased with geographic distance (Figure 4A), consistent with isolation by distance. Poisson regression models across all length bins indicate that geographic distance, regions and the interaction between geographic distance and regions are statistically significant (*p* <10^-16) predictors of the mean number identity by descent segments. A likelihood ratio test and model selection with AIC both had significant region by distance interactions, indicating regions differed in the rate of distance decay in haplotype sharing (likelihood ratio test across all bins, p <10^-16; AIC: 1-3 cM, model1 with interaction: 4.7365e+08, model2 without interaction: 4.7674e+08; 3-5 cM model1: 6.6756e+08, model2: 6.7161e+08; >5 cM model1: 8.1954e+08, model2: 8.2428e+08). The distance decay within haplotype sharing was particularly strong in East Africa despite having the smallest geographic extent. This pattern may reflect reduced effective migration rates or reduced gene flow between highland and lowland landraces in Ethiopia primarily aided by adaptation to higher altitudes as described in a recent pangenome study (Morris et al., 2025).

**Figure 4.**
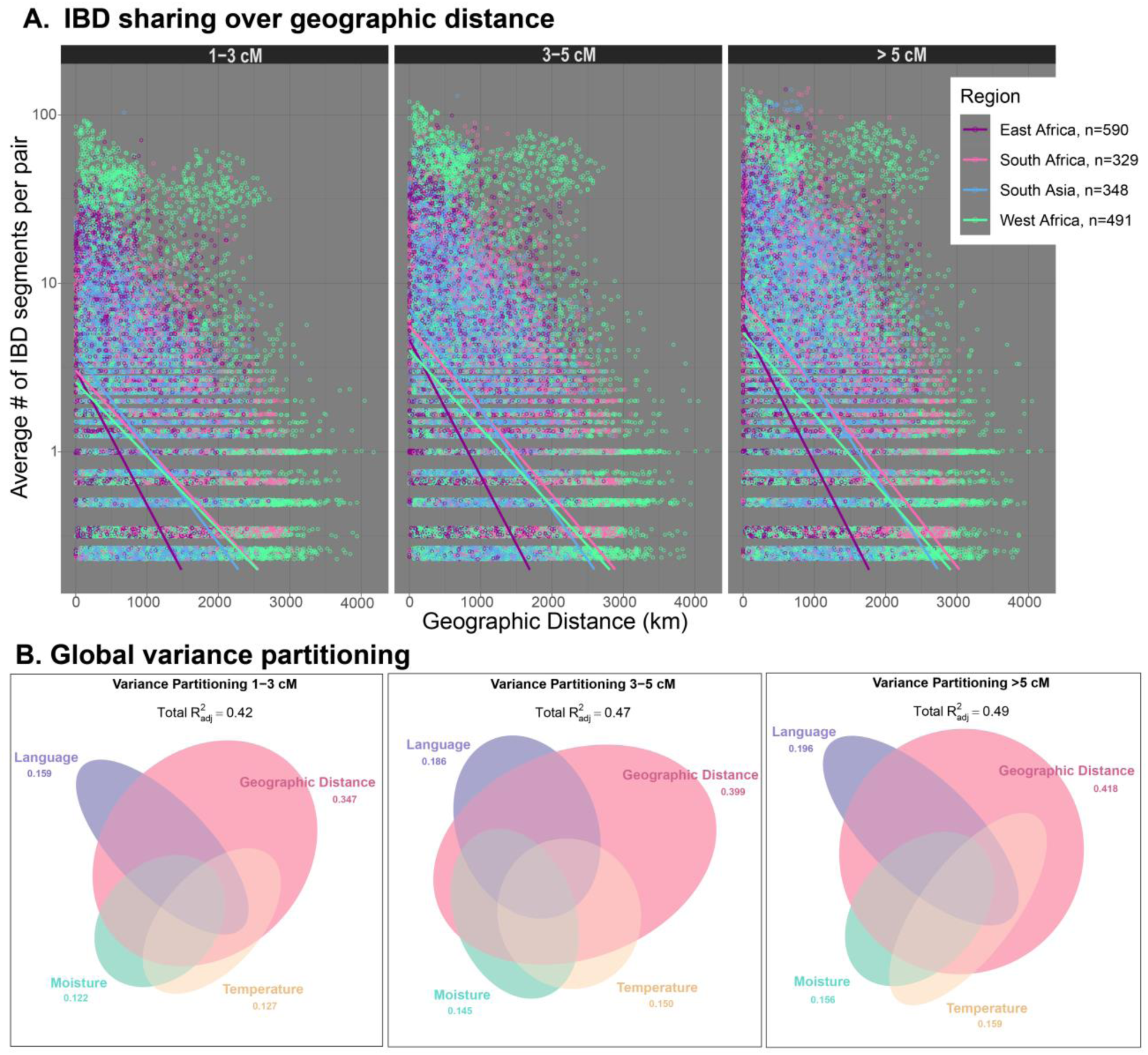
A) Average number of shared IBD tracts (log scale) across haplotype pairs (excluding pairs with no shared ancestry) and geographic distance within multiple identity by descent length bins, representing different timescales. The 1-3 cM bin represents older relationships, 3-5 cM intermediate and >5 cM recent relationships. The trend lines correspond to predicted weighted poisson regression values for each region across all pairs (weight by sample size). **B)** Variance partitioning for predictor variables moisture, temperature, geographic distance, language across identity by descent length bins. The adjusted R^2^ value by each predictor variable represents the degree of shared ancestry explained solely by that predictor. Ellipse size corresponds to the proportion of variation explained by variables in each category, with the ellipse overlap indicating collinear variation in predictors that explained variation in shared ancestry.

Distance decay in haplotype sharing was strongest for >5 cM segments followed by 3-5 cM and 1-3 cM as the degree of shared ancestry (i.e the average number of IBD segments per pair) increased over time. Increased geographic decay among recent relationships relative to older relationships is consistent with migration models where the geographic distance from an individual’s genetic ancestor is expected to increase at a rate of √t, looking backward in time (Kelleher et al., 2016; Ralph & Coop, 2013). In other words, genotypes are more likely to have genetic ancestors in close geographic proximity in recent times than in the distant past.

### Multicollinearity among geography, language, and climate explains sorghum genetic variation

To assess the extent to which geographic distance (or travel time), language and climate explained genetic variation in landraces we used variance partitioning with redundancy analysis. We found most SNP variation was individually explained by geographic distance and language (R^2^_a**dj**_ = 0.223 and 0.107, respectively) followed by temperature and moisture (R^2^_ad**j**_ = 0.088 and 0.082, respectively) (Figure 5). Together, the climatic variables explained 12% of genetic variation while geographic space collinear with language explained 24% of SNP variation.

**Figure 5.**
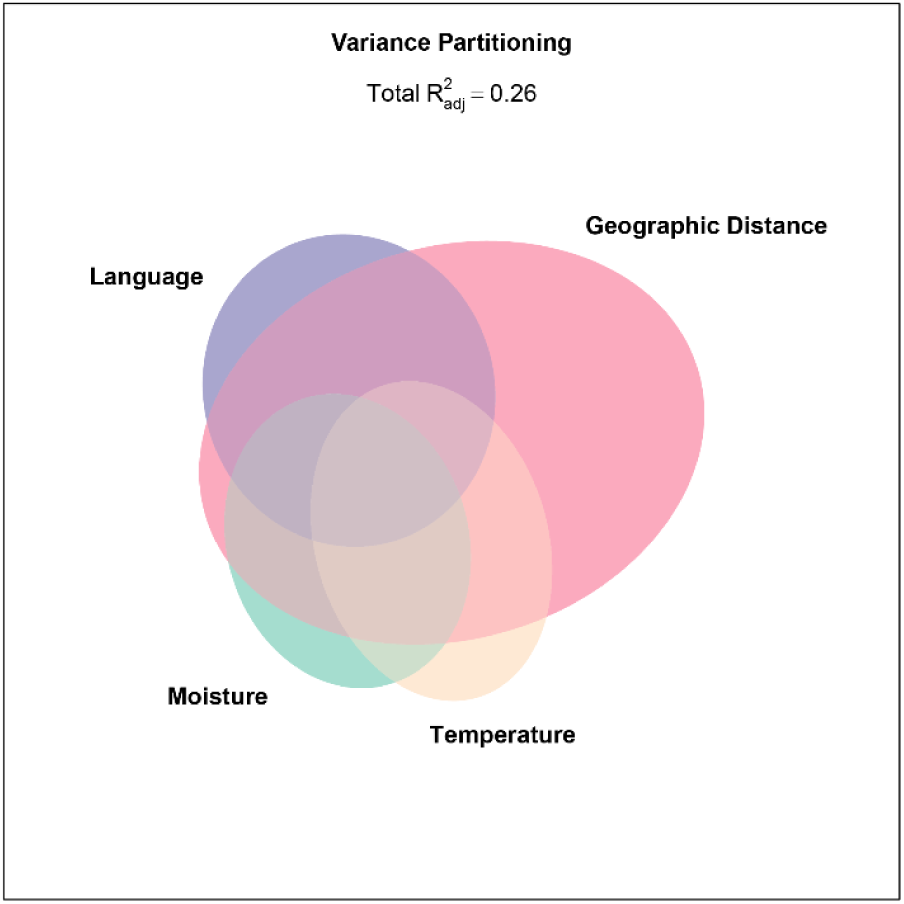
Variance partitioning for predictor variables moisture, temperature, geographic distance, language using SNPs as the multivariate response factor. Ellipse size corresponds to the proportion of variation explained by variables in each category, with the ellipse overlap indicating collinear variation in predictors that explained variation in SNPs.

Temperature collinear with geographic distance as well as moisture collinear with geographic distance explained 23% of SNP variation. Using travel time in place of geographic distance gave similar results: travel time and language (R^2^_a**dj**_ = 0.228 and 0.107, respectively) individually explained the most genetic variation in sorghum landraces followed by temperature and moisture (R^2^_adj_ = 0.088 and 0.082, respectively) (S3). Most SNP variation (25%) was explained by travel time collinear with language followed by moisture collinear with travel time (R^2^_adj_ = 0.248) and temperature collinear with travel time (R^2^_adj_ = 0.245). Moisture collinear with temperature explained 12% of SNP variation. Comparatively, the travel time model explained 27% of genetic variation in sorghum landraces while geographic distance model explained 26%.

When we considered the relationship between genomic variation in these predictors across time, we found the relative explanatory power of individual and combined factors increased for more recent relationships, with 42%, 47% and 49% of shared ancestry explained by predictors for the 1-3, 3-5 and >5 cM bins respectively (Figure 4B). For simplification, we will use 1, 3 and 5 to represent 1-3, 3-5 and >5 cM bins below. Similar to our variance partitioning analysis utilizing SNPs as the multivariate response factor (Figure 5), geographic distance (1: 34%, 3: 39%, 5: 41%) followed by language (**1:** 15%, **3:** 18%, **5:** 19%) independently explained the most shared ancestry across all timescales. By contrast, moisture and temperature equally explained the degree of shared ancestry over time (**1:** 12%, **5:** 15%) with the exception of the intermediate timescale where temperature and moisture explained 15% and 14% of shared ancestry respectively. With the exception of the oldest relationships (1-3 cM segments), we observed an equal degree of collinearity between geographic distance and language as well as geographic distance and moisture over time (**3:** 43%, **5:** 45%). When we compared the relative number of differences (adjusted r-squared values) among individual factors within each bin, we observed the highest gap in the 3-5 cM bin (56) followed by > 5 cM (52), and 1-3 cM (44).

## Discussion

Understanding the patterns of geographic structuring and genetic relatedness across space and time as well as the relative contribution of factors to genetic diversity may allow us to uncover sources of genetic variation, which may help the conservation of diversity for improvement of staple crops globally. Presently, crop-related research studies have focused on reconstructing the origin and dispersal patterns of staple crops as well as the molecular, genetic and environmental mechanisms contributing to plant resilience and adaptation (Alam et al., 2021; Caproni et al., 2023; Silva et al., 2015; Zhang et al., 2021). Geographic space and climatic factors like temperature and precipitation in particular have been utilized to model gene flow patterns and explain genetic differentiation within and between populations (Gutaker et al., 2020; Lasky et al., 2015; Olatoye et al., 2018). Nevertheless, the factors that explain genetic variation outside of climate or space as well as the elements within climate and space that contribute to fine-scale population structure have received little attention until recently. Human-mediated gene flow, represented as human movement over landscapes, as well as cultural factors like language have gained recognition as contributors to crop genetic structure, contextualizing historical diffusion of crops across space (Benz et al., 2007; Gilabert et al., 2025; Gutaker et al., 2020; Samberg et al., 2013). However, few studies have resolved the degree to which geographic space, environment, human movement and culture conjointly and independently explain genetic variation. And despite the importance of geographic space in modeling genetic relationships, it has been difficult to discern which relationships are more recent, or how the pattern of shared ancestry changes across geographic space. This study expands upon previous findings through an integrative examination of broad and fine-scale patterns of population structure to unveil the population genomic history of sorghum. By modeling intercontinental, regional and linguistic patterns of geographic structure using paired landraces, we build upon analyses of effective migration surfaces to reveal a strong pattern of isolation by distance at a large-scale, which when stratified by region and language families reveals further structuring. By employing identity by descent segments, we observed how the pattern of genetic relatedness across space changed over time and across regions, which expands upon models of early diffusion.

### Isolation by distance and travel time explain sorghum diversity and distribution

The spatial distribution of genomic variation in sorghum landraces across Africa and Eurasia (Figure 1A & B) presents a pattern of isolation by distance (Wright, 1943) and travel time, where gene flow between landraces is reduced as geographic distance or travel time increase. Globally, the plateau in genetic distance at ∼10,000km may reflect the rare occurrence of long-distance dispersal events between populations (Ibrahim et al., 1996; Jordano, 2017). Contrary to our expectations, globally the geographic distance model was a better fit to sorghum genomic variation than was estimated travel time. By contrast, Gutaker et al. (2020) found that travel time was a better fit than geographic distance for *japonica* rice landraces (though not *indica* rice).

However, when we compared regional models, we found the travel time model was a better fit in East Africa, West Africa and South Asia while geographic distance was a better fit in South Africa. This result suggests that our travel time model may capture relative variation in gene flow at the within-region level, and that this model performed better at modeling shorter travel routes. While improvements could be implemented in our travel time model, other biological or cultural factors like variety-specificity and/or differences in geographic constraint among linguistic groups may explain why there are scale-dependent differences in model fit. Limitations for the travel time model could be related to the implementation of waypoints and more specifically the exclusion of waypoints in South Asia, which may have better represented human movement between continents. Future modifications may involve a systematic division of conductance layers into demes and the inclusion of a desert-specific travel model.

### Sorghum genetic structure explained by hierarchical linguistic differences

We found evidence of linguistic structuring within language families and individual languages. Language families, which may be a proxy for cultural preferences and cultivation practices as well as gene flow mediated by shared culture, explained up to 4% of genetic variation in both our estimate of hierarchical F_ST_ and multicontinental F_ST_. Individual languages explain 13% of total genetic variation and 10% of total SNP variation within language families (Figure 3A).

Westengen et al. (2014) and more recently, Gilabert et al. (2025) provided systematic evidence for ethnolinguistic population structure, but only within Africa. Linguistic structuring of sorghum landraces in Africa was observed among language families and subclasses (Gilabert et al., 2025). Our analysis of hierarchical F_ST_ corroborates these findings across multiple continents and language families. In other words, a majority of SNP variation explained by language families can also be found within individual languages but there is also a small portion within language families that only individual languages can explain, which suggests that individual languages explain more genetic diversity than language families. This difference may be attributable to differences in granularity between individual languages and language families. For example, a large geographically widespread language family like Atlantic-Congo has a diversity of individual languages which in themselves may exhibit differing patterns of spatial structuring (i.e variation in the degree of genetic differentiation based on geographic proximity). Thus, the SNP variation explained by language families, particularly for larger language families with a large geographic extent, may not fully capture fine-scale population structure.

Like population genomic similarity, researchers have also found that language variation is spatially autocorrelated (Barbieri et al., 2022). Linguistic spatial autocorrelation may thus drive differences in geographic structure across language families due to selection and shared gene flow of sorghum driven by human groups sharing language. Despite the simplicity of language family groupings in analyses of population structure, they can still be utilized to assess differing patterns of spatial structuring at a global scale. Our analysis of spatial structuring across the language families with the largest sample sizes showed that the amount of genetic diversity, geographic extent and distribution across space differs despite the consistent isolation by distance pattern (Figure 3B). By contrast, sorghum landraces attributed to the Mande language family exhibited evidence of genetic divergence, with two distinct genetic clusters; this division may be a result of reproductive isolation between the highly diverged *guinea margaritiferum* variety and more common, *guinea* and *durra* variety in west Africa, which we elaborate on below. Ultimately these analyses show that language can be a predictor for sorghum genetic structure and that language families in themselves exhibit differing patterns of isolation by distance. While we only considered language as a proxy for culture, other factors like ethnicity (Faye et al., 2019), and farmer social dynamics may play a role (Leclerc & d’Eeckenbrugge, 2012; McGuire, 2007). Future projects may consider implementing these factors to represent cultural preference at any geographic scale.

### Spatiotemporal differences in genetic and geographic structure

Although landraces exhibited a pattern of isolation by distance and travel time across all regions, we observed considerable variation in the distribution of landraces across space (Figure 1C), which suggests there are regional differences in the pattern of isolation by distance. Gilabert et al. (2025) and Morris et al. (2013) identified a genetically divergent clade within West Africa associated with the *guinea margaritiferum* variety, which may have been domesticated independently or carry introgression from wild sorghum. Our assessment of regional and linguistic differences in spatial structuring was consistent with these findings (Figure 1C & 3B). Among the considered subregions, a subset of West African landraces exhibited a distinct genetic division, which may be evidence of reproductive isolation. The sorghum landraces assigned to the Mande language family (Figure 3B), which was restricted to landraces within pockets of West Africa such as Southwest Mali and Sierra Leone (Figure 3C), were highly divergent in comparison to other paired landraces in West Africa, South Africa and South Asia. The presence of two divergent sorghum genetic clusters extended past the geographic range of the Mande language family, which suggests that this variety has expanded across communities with distinct language families within the region. South Asian landraces also exhibited a degree of genetic divergence, which may follow the East-West division of the Indo-European and Dravidian language families, and associated *durra* from drier desert climates and *guinea* varieties prevalent in the relatively wetter tropical savanna regions (Figure 3B & C) (Lasky et al., 2015; Morris et al., 2013). Genetic divergence between South Asian landraces, specifically in India, may not be as prominent because of interbreeding between the prevalent *durra* and *guinea* varieties in areas where they co-occur (Morris et al., 2013). Comparatively, the distribution of landraces in East Africa reflects large-scale isolation by distance pattern, which may be indicative of increased opportunity for admixture over time. As a hotspot for sorghum diversity, admixture within East Africa may have been bolstered by cultural exchange (e.g., seed sharing and emigration) (Menamo et al., 2021), which could have impacted the degree of genetic differentiation.

In our assessment of temporal differences in spatial structuring, we observed regional differences in the relationship between geographic proximity and shared ancestry over time (Figure 4A). Our findings suggest that landraces that are geographically proximate are more likely to share genetic ancestry than those that are more distant. However, the rate of geographic decay varied regionally and across different timescales. We observed the steepest geographic decay among East African landraces, where there was a sharp decline in the degree of shared ancestry as geographic distance increased. This pattern may be attributable to differences genetic structure and adaptation along altitudinal gradients in Ethiopia as found in previous studies in sorghum (Gilabert et al., 2025; Morris et al., 2025) as well as in barley (Hadado et al., 2010). The conjoint effects of altitude and differential farmer practices also exhibited strong genetic differentiation in Ethiopian barley (Samberg et al., 2013), which could play a role in this context. By contrast, many West African landraces presented a notable pattern of high shared ancestry for many pairs of landraces at larger distances, which may be attributable to rapid spread of some sorghum genotypes across long latitudinal belts of similar climate in this region (Morris et al., 2013). Over time with longer chromosomal segments, the degree of shared ancestry increased across all regions, particularly in the >5 cM bin, a pattern that is expected under limited migration models where distance from an ancestor increases backwards through time (Kelleher et al., 2016; Ralph & Coop, 2013). Greater shared ancestry at closer distances in recent times may have also been a result of limited dispersal, local adaptation to climates and differential preferences among communities, which have likely strengthened the pattern of isolation by distance among more recent relationships. Regional differences in the pattern of genetic relatedness across space and time may further contextualize the regional differences in the pattern of spatial structuring we observed (Figure 1C).

Our variance partitioning analysis showed that the relative explanatory power of geographic distance, climate and language as well as their conjoint effects increased when comparing larger haplotypes representing more recent relationships (Figure 4B). Consistent with our assessment of the effect of these predictors on SNP variation (Figure 5), we found that geographic distance and language consistently explained the most identity by descent sharing over time.

The explanatory power of individual and conjoint factors was relatively similar over time, suggesting the processes determining large scale population structure have been largely consistent through time. However, the degree of shared ancestry explained by temperature was slightly higher than moisture in the intermediate timescale. Similarly, geography collinear with language was higher than geography collinear with moisture in the earliest timescale (1-3 cM). This suggests that there are some slight differences in the importance of variables over time. For example, in the earliest timescale, geography collinear with culture may have been a more important contributor to shared ancestry than geography collinear with moisture due to limited dispersal favoring communities sharing the same language family, and reduced gene flow between different climates in the early diffusion period of sorghum landraces.

### Multicollinearity of spatial and human-related factors explain most genetic variation

Our variance partitioning analysis suggests that geographic space and travel time individually explained the most genetic variation in sorghum landraces (Figure 5). However, 24% and 25% collinearity with language for space and travel time respectively indicates that detailed study of human history and preferences may be required to better disentangle culture from isolation by distance to further contextualize crop spatial dynamics. We also observed 24% collinearity between geographic distance and the climatic factors moisture and temperature, which aligns with findings in previous studies that consider space and environment (Faye et al., 2019; Lasky et al., 2015; Olatoye et al., 2018). These findings suggest that SNP variation in multicontinental sorghum landraces can be explained by geography and climate as established regionally in Nigerian (Olatoye et al., 2018) and Senegalese (Faye et al., 2019) sorghum, as well as globally (Lasky et al., 2015). Collinearity between geographic distance and climate may be partially explained by a pattern of isolation by distance and spatial autocorrelation in climate conditions. Here we found that geographic distance and language explained more genetic variation than climate alone or space and climate, highlighting the importance of components of human culture like language in sorghum population structure.

## Conclusion

Climate change in certain regions may limit the variety of crops that can be cultivated, making drought-tolerant crops like sorghum essential for future agriculture (Rosenow et al., 1983; Tuinstra et al., 1996). Understanding the factors that influence dispersal patterns of crops may allow us to make predictions about the adaptive capacity and survival probability of staple crops under temporal environmental and habitat variability. Furthermore, the patterns identified here help place in context knowledge developed about the history of specific adaptive quantitative trait loci (Morris et al. 2025). Our assessment of the ecological and environmental factors that affect sorghum diversity suggests that spatial proximity, human movement, culture, and climate differentially impact genetic structure relative to each other as well as over space and time. The pattern of isolation by distance and travel time across Africa, Eurasia and South Asia indicates that gene flow between landraces is limited by geographic distance and human-mediated dispersal, particularly between more distant varieties. We also observed considerable variation in spatial structuring across language families and regions and time, which highlights the importance of these factors in determining genetic variation. Crop breeding and/or conservation efforts may consider the effects of culture and language as partial predictors for genetic diversity.

Additionally, this study may encourage researchers to explore beyond language families, as more genetic variation may lie within languages alone. By leveraging nearly 2000 GBS georeferenced sorghum landraces across a large geographic range and associated climatic data we were able to assess population structure across different spatial and temporal scales, highlighting the utility of agricultural landscape genomics (Campbell et al., 2025). Ultimately, this research may provide a historical framework for sorghum geneticists and further understanding of evolutionary drivers of sorghum diversity.

## Acknowledgements

Travel time calculations were run using Bridges-2 resources from the Pittsburgh Supercomputing Center through allocation BIO230186 from the Advanced Cyberinfrastructure Coordination Ecosystem: Services & Support (ACCESS) program, which is supported by National Science Foundation grants #2138259, #2138286, #2138307, #2137603, and #2138296.

This work was supported by the National Science Foundation Graduate Research Fellowship Program under Grant No. DGE1255832 to EEVC, Gates Foundation grant “Mining useful alleles for climate change adaptation from CGIAR gene banks (INV-030574)” to GPM and JRL, and by National Institutes of Health grant R35GM138300 to JRL.

