## Supplementary Materials for "Population genomic structure of sorghum landraces across landscape, environment and culture"

### Supplemental Materials

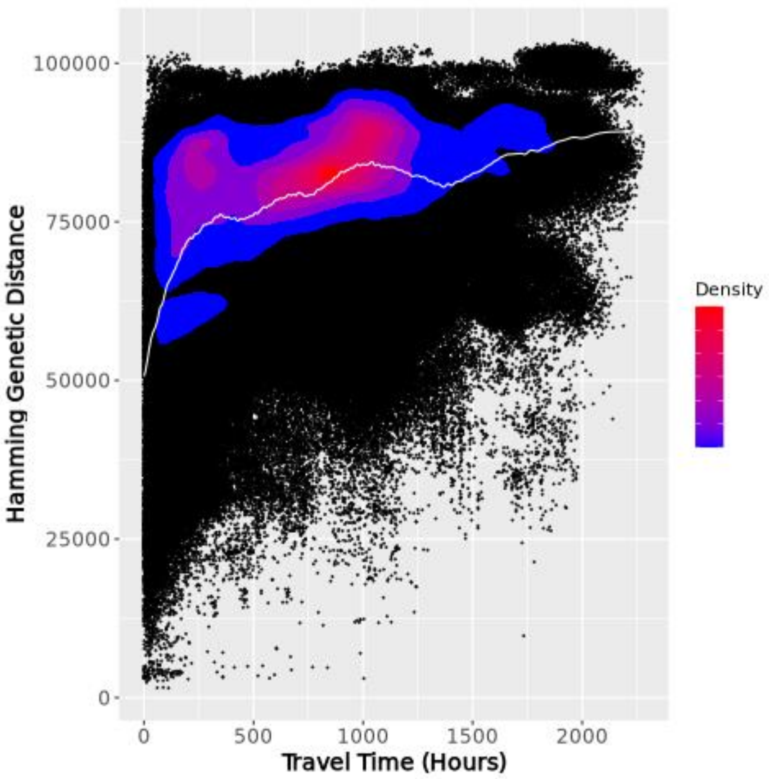


S1. Scatterplot of travel time and genetic distance 1,806 landraces. Spline (white line) added to show distribution pattern. $R^2$=0.188

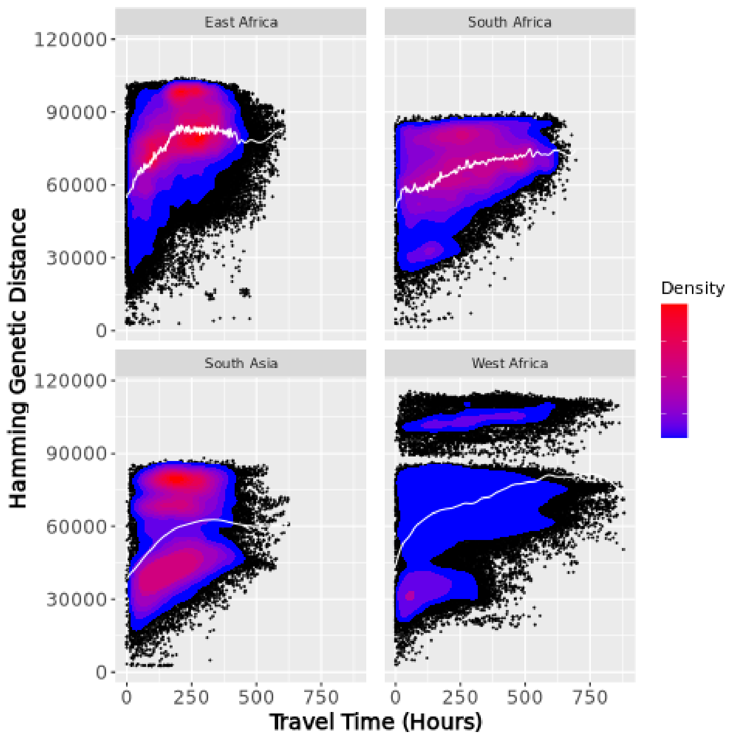


S2. Scatterplot of travel time in subregions East Africa ($R^2$= 0.140), South Africa ($R^2$= 0.131), and West Africa ($R^2$= 0.111), South Asia ($R^2$= 0.078). Spline (white line) added to show distribution pattern.

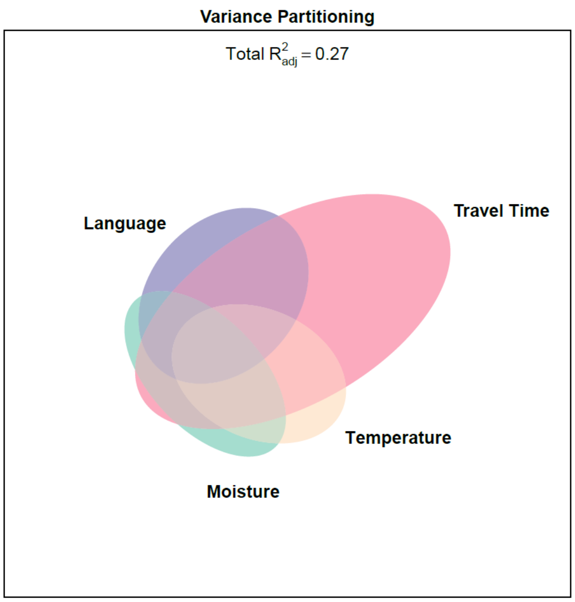


S3. Variance partitioning for predictor variables moisture, temperature, travel time, language using SNPs as the multivariate response factor. Ellipse size corresponds to the proportion of variation explained by variables in each category, with the ellipse overlap indicating collinear variation in predictors that explained variation in SNPs.

$R^2_{adi}$ = 0.27. Residuals = 0.72

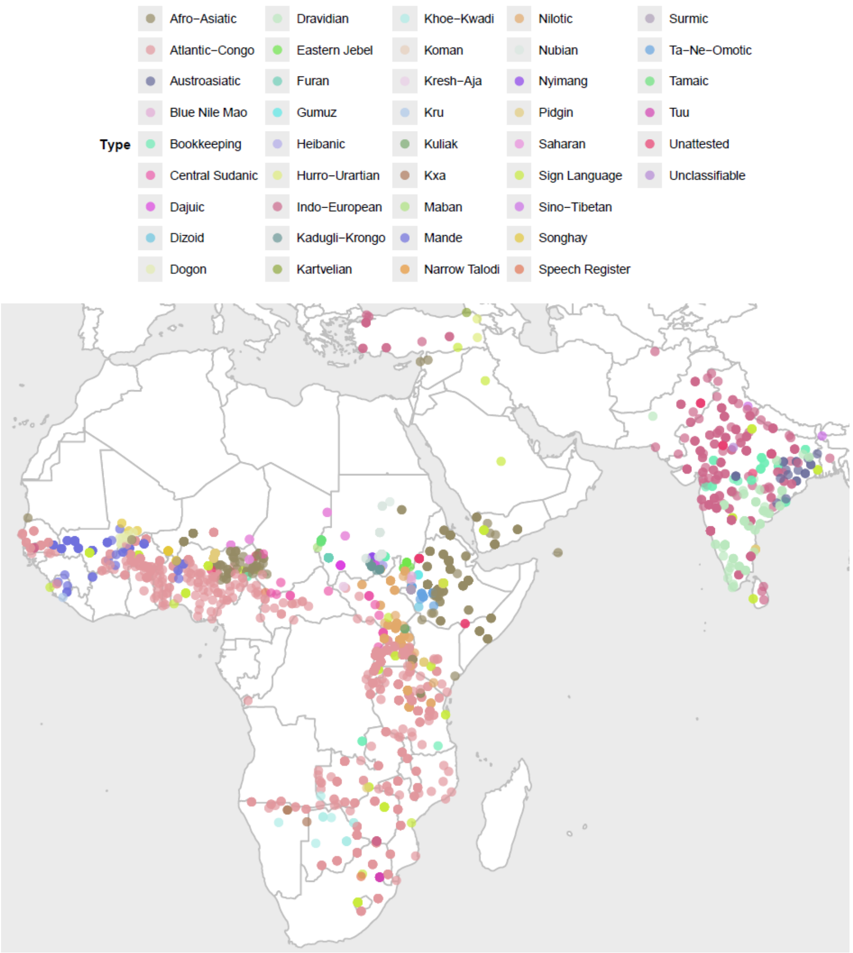


S4. Map of 1806 landraces and associated 42 unique language families.

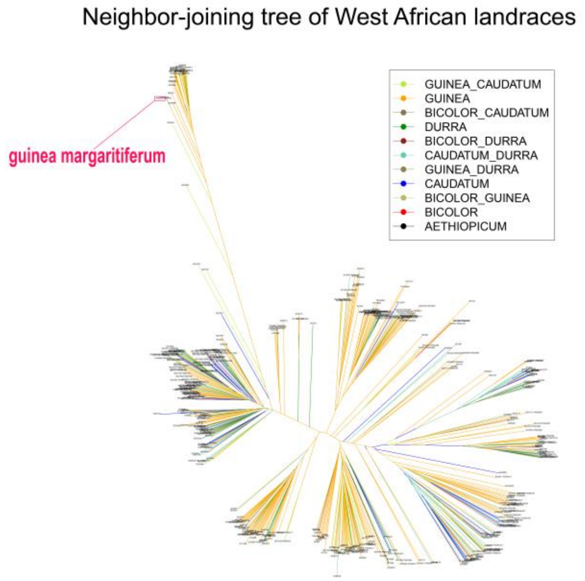


S5. Neighbor-joining tree depicting genetic relatedness (Hamming distance) among West African landraces. Legend colors denote variety names. The landrace "IS3620," which is categorized as the *guinea margaritiferum* subrace, is highlighted.

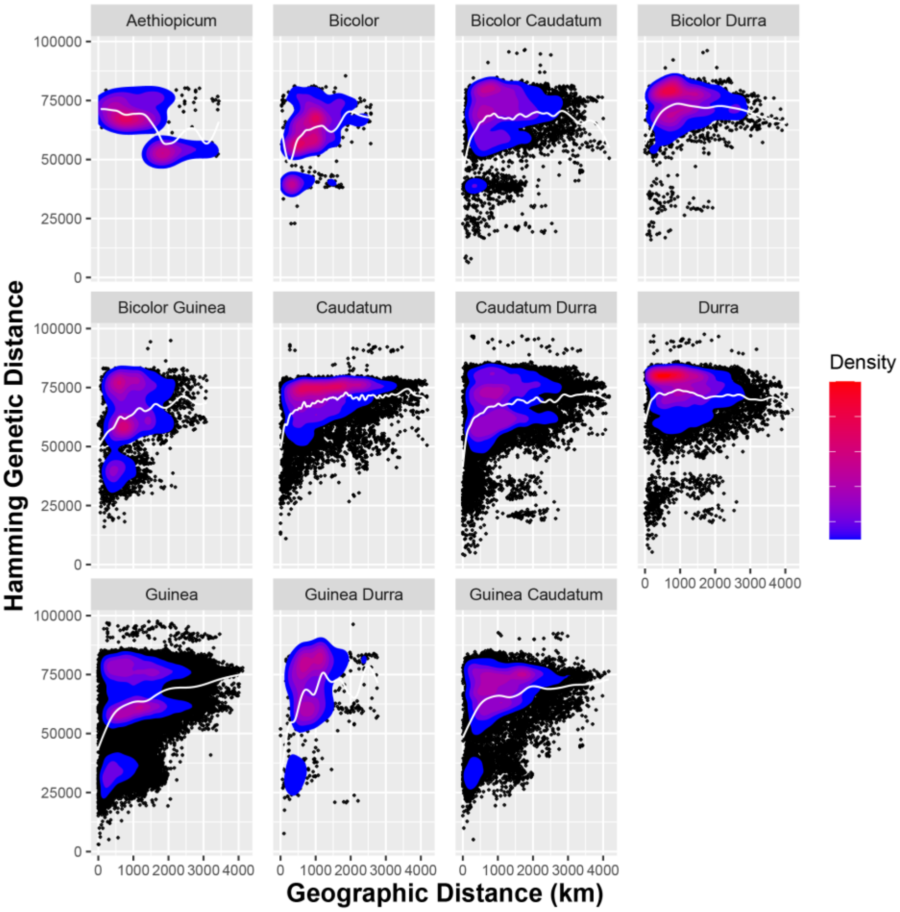


S6. Scatterplot of geographic distance and genetic distance for 11 varieties present among 491 West African landraces (diverged *guinea margaritiferum* varieties removed). The 30 *margariteferum* landraces removed were identified using hclust (k=2) dendogram. Spline (white line) added to show distribution pattern.

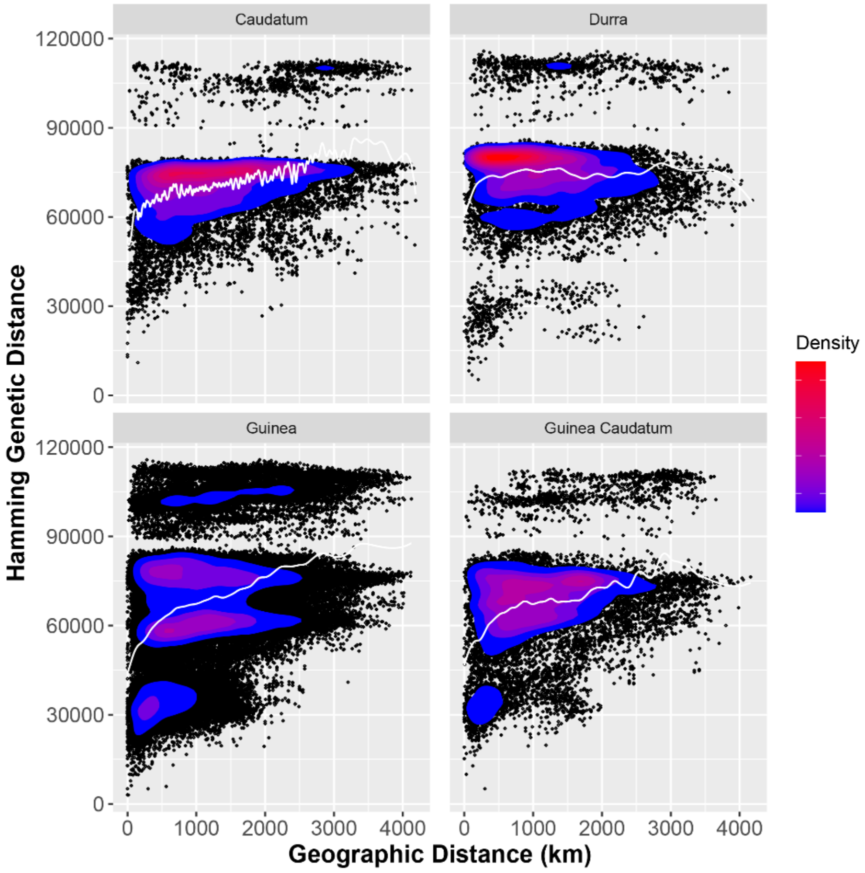


S7. Scatterplot of geographic distance and genetic distance for 4 varieties present in West Africa. The distribution pattern of the Guinea variety reflects the same divergence pattern present in West Africa. Spline (white line) added to show distribution pattern.
